# The Origins and Consequences of Localized and Global Somatic Hypermutation

**DOI:** 10.1101/287839

**Authors:** Fouad Yousif, Stephenie D. Prokopec, Ren X. Sun, Fan Fan, Christopher M. Lalansingh, Erik Drysdale, David H. Park, Lesia Szyca, PCAWG Network, Paul C. Boutros

**Affiliations:** Ontario Institute for Cancer Research, Toronto, Canada; Department of Pharmacology & Toxicology, University of Toronto, Toronto, Canada; Department of Medical Biophysics, University of Toronto, Toronto, Canada

## Abstract

Cancer is a disease of the genome, but the dramatic inter-patient variability in mutation number is poorly understood. Tumours of the same type can differ dramatically in their mutation rate. To improve our understanding of potential drivers and the consequences of the underlying heterogeneity in mutation rate across tumours, we evaluated both local and global measures of mutation density (both single-stranded and double-stranded DNA breaks) in 2,460 tumours across 38 cancer types. We find that SCNAs in thousands of genes are associated with elevated rates of point-mutations, while conversely, SNVs in dozens of genes are associated with specific patterns of DNA double-stranded breaks. To supplement this understanding of global mutation density, we developed and validated a tool called SeqKat to identify localized regions of hypermutation (also known as kataegis). We show that rates of kataegis differ by four orders of magnitude across tumour types and that tumours with *TP53* point mutations were 2.6-times more likely to harbour a kataegic event than those without. Furthermore, we identify novel subtypes of kataegic events not associated with aberrant APOBEC activity and found that kataegic events were associated with patient survival in some, but not all tumour types. Taken together, we reveal a landscape of genes driving localized and tumour-specific hyper-mutation, and reveal novel mutational processes at play in specific tumour types.

## Introduction

Genome instability is one of the hallmarks of cancer, and cancer is often referred to as a “disease of the genome”^1^. Just as cancers are heterogeneous over time and space^2-4^, they are also heterogeneous in the number of mutations they harbour - their “mutational density”. Some tumour types, like melanomas and lung cancers, harbour hundreds of thousands or even millions of single nucleotide variants (SNVs). These large mutational burdens are thought to reflect the effects of environmental carcinogens, like UV radiation and the by-products of cigarette smoke. Other tumour types, like prostate cancers, can harbour only a few hundred SNVs^5-11^. This variability in SNV mutational density across tumour types has been well-demonstrated in previous pan-cancer exome sequencing studies^12,13^.

It is less clear however, what drives some tumours to harbour more DNA damage than others. Even within an individual cancer type, individual tumours of similar clinical grade and stage can vary by orders of magnitude in their number of SNVs. Theoretical modeling studies have attributed large fractions of the divergence across tumour types to differences in the number and rate of replication-induced errors, rather than the effects of heredity or environmental influences^14-16^. What drives these differences in mutational density at the SNV level? Are these differences in mutational density in point-mutations reflected by similar trends in somatic copy number aberrations (SCNAs), translocations and other copy-neutral structural variants (SVs) and in localized hypermutation events such as kataegis?

To address these questions, we evaluated both local and global measures of mutation density: including numerous metrics of both single-stranded and double-stranded DNA breaks. Using mutation data from the PCAWG Network for 2,460 tumours across 38 cancer types, we have uncovered potential mechanisms and/or consequences associated with mutational density. Specifically, we have identified individual genes whose mutation status is associated with changes in the mutation density of different types of aberrations. For example, we identify SNVs in dozens of genes as associated with changes in the burden of somatic copy number aberrations (SCNAs) and the number of copy-neutral SVs in a pan-cancer analysis. These candidate drivers of mutation density preferentially occur early in tumour evolution, appearing clonally in all cells of a tumour.

To further expand this understanding to localized mutation density, we developed and validated SeqKat, a tool to identify genomics regions with statistically significance increases in local point-mutation frequency. These regions are then classified as either kataegic or non-kataegic hypermutation events, based on mutation type and trinucleotide context. Rates of kataegis differed dramatically across tumour types, with malignant lymphomas having a particularly high rate. Furthermore, we identify novel subtypes of kataegic events not associated with aberrant APOBEC activity, and find that these are localized to specific genomic regions and enriched for MYC-target genes. Kataegic events were associated with patient survival in some, but not all tumour types, highlighting a combination of global and tumour-type specific effects. Taken together, we reveal a landscape of genes driving localized and tumour-specific hyper-mutation, and reveal novel mutational processes at play in specific tumour types.

## Results

### Experimental Design

The ICGC Pan-Cancer Analysis of Whole Genomes (PCAWG) analyzed the whole genomes sequences (WGS) of 2,703 tumour-normal pairs of 37 histological subtypes using a consistent bioinformatics pipeline (https://dcc.icgc.org/pcawg#!%2Fmutations)^17^. We excluded samples flagged for poor quality, which represented additional specimens from a single donor, where donor sex was unknown or where donor age was unknown. The remaining 2,460 tumours were evaluated for density of somatic single-stranded breaks, double-stranded breaks, hypermutation and kataegis (**Supplementary Figure 1; Supplementary Table 1**). Selection procedures for tumours focused on larger, surgically-managed tumours that yielded sufficient DNA for sequencing: the PCAWG marker paper outlines variant calling, coverage and other aspects^17-19^.

### Drivers of Single Nucleotide Variation

We first examined the density of somatic SNVs across the PCAWG cohort, and observed a broadly consistent number of SNVs, including both coding and non-coding variants, across the genome (excluding chromosome Y), with a median of 12,950 SNVs within each 1 Mbp bin (**Figure 1A**). Regions with few SNVs largely correspond to centromeres and telomeres, and after control for differential coverage across samples, hypermutated bins did not encompass known driver genes (**Supplementary Figure 2A**). Rather, these reflect replication-timing effects, environmental exposures and other signals^8,20,21^. Across tumour types, we confirm the relatively small intra-tumoural heterogeneity seen in previous studies (Figure 1B)^6,22-25^ with 99.3% of variance in SNV mutational burden between tumour types and only 0.7% of variance within them. However there are also significant differences between tumour types in their variability in SNV mutation density. For example at one extreme, median SNV mutation density of a melanoma was 26.4 SNVs/Mbp, but this varied dramatically (SD = 54.25, IQR = 45.59). By contrast in pilocytic astrocytomas, mutation density was both lower (0.06 SNVs/Mbp) and much more consistent across tumours (SD = 0.08, IQR = 0.076; **Supplementary Table 2**). Thus not only does mutation density vary significantly between tumour types, but so too does its consistency within them.

**Figure 1:**
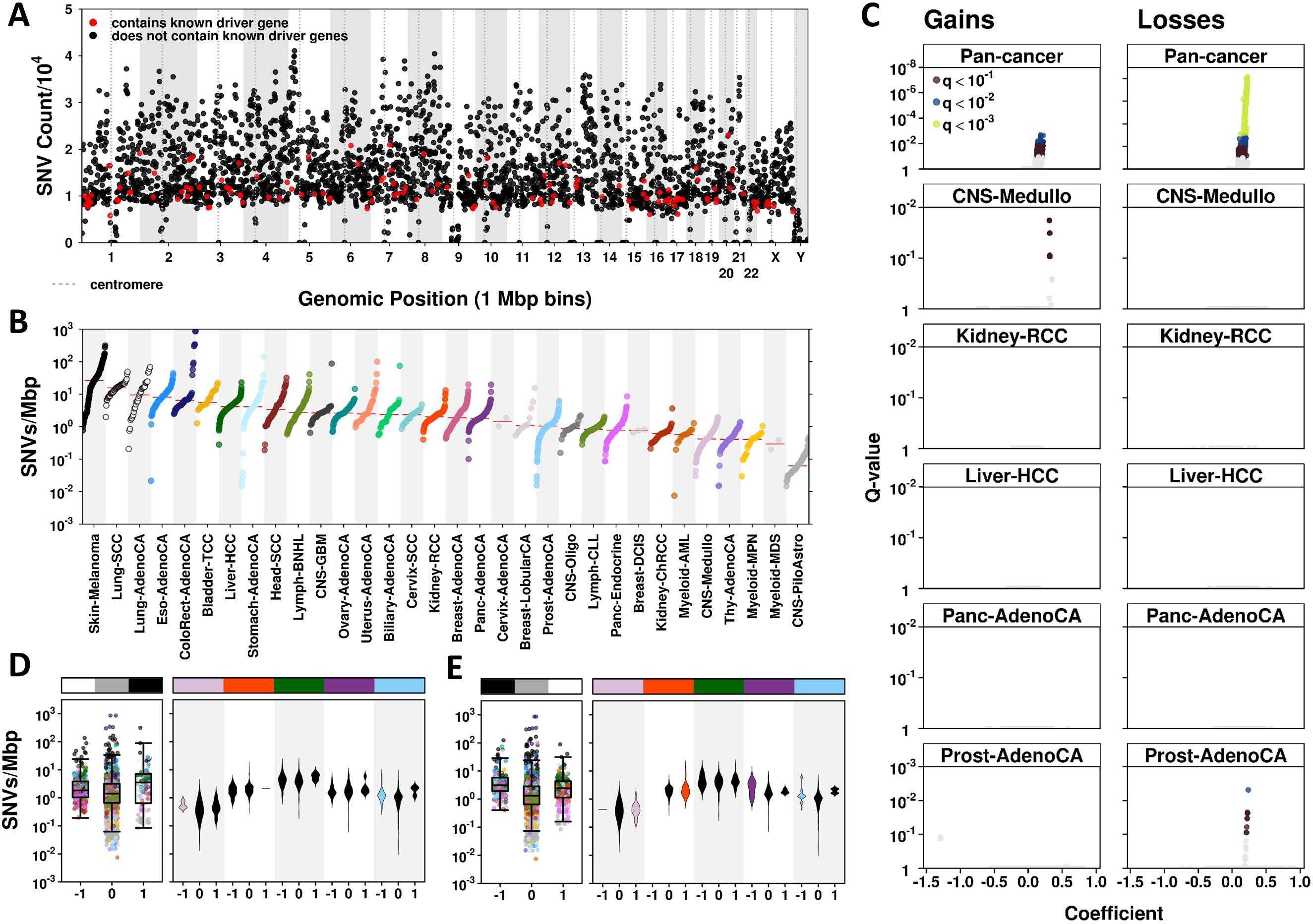
Specific SCNAs are associated with an increased SNV density. A) Total SNV count across all available patients within each 1Mbp bin along the genome; red points indicate bins containing known driver genes. B) Mutation rate (SNVs/Mbp), stratified by tumour type; red bars indicate median SNVs/Mbp for each type. Volcano plots showing the coefficient and FDR-adjusted p-value for all C) CN gains or losses in the pan-cancer or individual tumour type models. D) Tumours with CN-amplification of *BCKDH8* show elevated SNV mutation rates in a pan-cancer multivariate analysis (left boxplot) and in multiple individual tumour types (violin plots stratified by CN type (deletion = −1, neutral = 0, amplification = 1); coloured violins reflect statistical mixed effects model, FDR < 0.1). The tumour types selected for subgroup analysis are those with the largest sample number (CNS-Medullo, RCC, HCC, pancreatic adenocarcinomas and prostate adenocarcinomas), in boxplots as indicated by the covariates with colours corresponding to part B. E) *CKMT2* loss is associated with elevated mutation density. Figure structure is similar to 1D).

To understand the drivers of increased somatic point mutation density, we focused on their rate per Mbp of covered sequence (SNVs/Mbp, **Supplementary Figure 2B**). We analyzed the relationship between SNVs/Mbp and somatic copy number aberrations (SCNAs) in 1,778 samples with both types of data available. Linear mixed effects modeling with variable intercepts for each tumour type was used to quantify the effects of individual SCNAs on SNV mutational density, controlling for confounding variables including patient sex, age and average tumour ploidy (since changes in ploidy changes the amount of DNA available to accumulate mutations; **Supplementary Figure 2C-E, Supplementary Table 3**). Tumour purity was not correlated with SNVs/Mbp (**Supplementary Figure 2F**) so was used only for validation of specific models.

Similarly, less than 1% of variance in this mutation metric could be attributed to sequencing site *(via* project-code), therefore this variable was not included in our models.

Our analysis identified 16 gene amplifications and 473 gene deletions significantly associated with increased SNVs/Mbp (**Figure 1C, top**; Bonferroni adjusted p-value < 0.01). No genes were associated with decreased SNV mutational density (**Supplementary Table 3**). Some genes were associated with very large changes in SNV mutational density, such as amplification of the nuclear-encoded mitochondrial gene *BCKDHB.* Tumours with this gain harboured an additional 1.66 SNVs/Mbp than those without it across the PCAWG cohort, corresponding to ~5,000 additional SNVs genome-wide (**Figure 1D, left panel**). This pan-cancer effect was independent of tumour purity. To confirm our multivariable modeling identified trends that occur in individual tumour types, we performed subgroup analyses on the five tumour types with the highest individual number of tumours in the PCAWG cohort: medulloblastoma, renal cell carcinoma (RCC), hepatocellular carcinoma (HCC), pancreatic and prostate adenocarcinomas. Here, tumour purity appeared to be a contributing factor (data not shown), as was the scarcity of *BCKDHB* amplifications in these subsets, with deletions showing a more significant association in medulloblastoma and prostate adenocarcinomas (**Figure 1D, right panel**).

Similarly, deletion of nearly all genes on the q arm of chromosome 5 were associated with an increase in SNVs/Mbp. This region encompasses the tumour-suppressor *APC*, which was deleted in 9.6% of samples, and on which we focused. Tumours with deletion of APC showed an increase of 1.64 SNV (**Figure 1E, left panel**); this tumour suppressor and is representative of the deletion event, associated with an increase of 1.64 SNVs/Mbp (~5,000 SNVs genome-wide). A further examination of the five tumour with the largest sample-number confirmed this trend with both pancreatic and prostate adenocarcinoma showing significant increases in SNVs/Mbp in tumours with *APC* deletions (**Figure 1E, right panel**). Thus individual gene-wise SCNAs, possibly representing larger SCNA segments, are associated with large-scale changes in somatic SNV mutational burden, both pan-cancer and within individual tumour types.

Genes with SCNAs associated with increased global somatic SNV mutational density were assessed for pathway enrichment. Amplified genes associated with increased SNVs/Mbp preferentially harboured GTPase activity (including *NET1* and *ECT2;* **Supplementary Table 4**). Conversely, deleted genes associated with increased SNV mutational density were enriched for interleukins (growth receptor binding and regulation of STAT) and subunits of the TFIIH holoenzyme (**Supplementary Table 4**). Amplifications were enriched on chromosome 10, while deletions were enriched on chromosome 5 (**Supplementary Table 5**). The majority of associated genes originated in subclonal tumour populations^26^. However, 251 genes affected by CN deletions associated with SNV mutational density were identified as clonal in at least 50% of patients (**Supplementary Table 6**), and these were preferentially located on the q arm of chromosome 5.

Finally, to better understand the tumour-type-specificity of candidate drivers of SNV mutational density, we performed genome-wide subgroup analyses on the five tumour types with the most samples: medulloblastoma, RCC, HCC, pancreatic and prostate adenocarcinomas. We replicated our mixed linear modeling strategy for each tumour-type independently, controlling for sex (where appropriate), age and tumour ploidy. Adjacent genes with identical copy-number profiles were collapsed within tumour types. Despite considerably smaller sample sizes (n ranging from 107-241), we identified multiple regions associated with increased SNV mutational density (Bonferroni < 0.1; **Figure 1C, Supplementary Table 7**). As before, effect-sizes could be very large; 9 unique CNA segments on chromosome 8 (representing 34 genes) carried CN deletions associated with increases in SNV mutational density of 1.63-1.73 SNVs/Mbp in prostate adenocarcinomas (**Supplementary Figure 3**). Similarly, 80 regions on chromosome 1 in medulloblastoma harboured CN gains (involving 452 genes) associated with an increase of 2.06-2.08 SNVs/Mbp (**Supplementary Figure 4**), independent of tumour purity. In both cases, findings were unique to a single tumour type and were not identified within the pan-cancer analyses. Taken together, these results uncover a landscape of pan-cancer and tumour-type-specific effects in driving changes in SNV mutational density. Indeed even the large PCAWG dataset employed here likely results in significant false-negative rates for smaller effect-size associations.

### Drivers of Copy Number Changes

We next sought to examine patterns of copy number gains and losses across tumour types. We considered a panel of copy-number summary features, including total SCNA count, proportion of the genome altered (PGA)^27^ and average SCNA length. These were further sub-categorized by the direction of change (gain, loss or overall; **Supplementary Table 1**).

We find that the well-known variability in SNV mutational density (**Figure 1A**) are paired to even larger intra- and inter-tumour type variability in their SCNA alteration patterns. For example, pilocytic astrocytomas had few SCNAs, predominantly amplifications (median PGA = 6.5%, **Figure 2A**). On the other end of the spectrum, chromophobe renal cell carcinoma (chRCC) was dominated by CN deletions, with few amplifications (median PGA = 0% and 37.3% for amplifications and deletions respectively; **Supplementary Figure 5, Supplementary Table 2**). Even within individual tumour types, these metrics ranged dramatically, with some having relatively balanced gains and losses and others showing large bias. For example, hepatocellular carcinoma (HCC) shows a balance of gains to losses (median ratio = 1), however with a large degree of variability around this (IQR = 2.58); similarly pancreatic adenocarcinomas show a median ratio of 0.55 (IQR = 0.79). This picture of large inter- and intra-tumour type heterogeneity was not restricted to any single feature of the CN landscape, but held true for many metrics of SCNA burden, both as an overall metric or when broken down to gains and losses separately, as well as for indirect features such as average ploidy and purity. In particular, dramatic variability was observed in the distributions of SCNA length across tumour types (**Supplementary Figure 6**).

**Figure 2:**
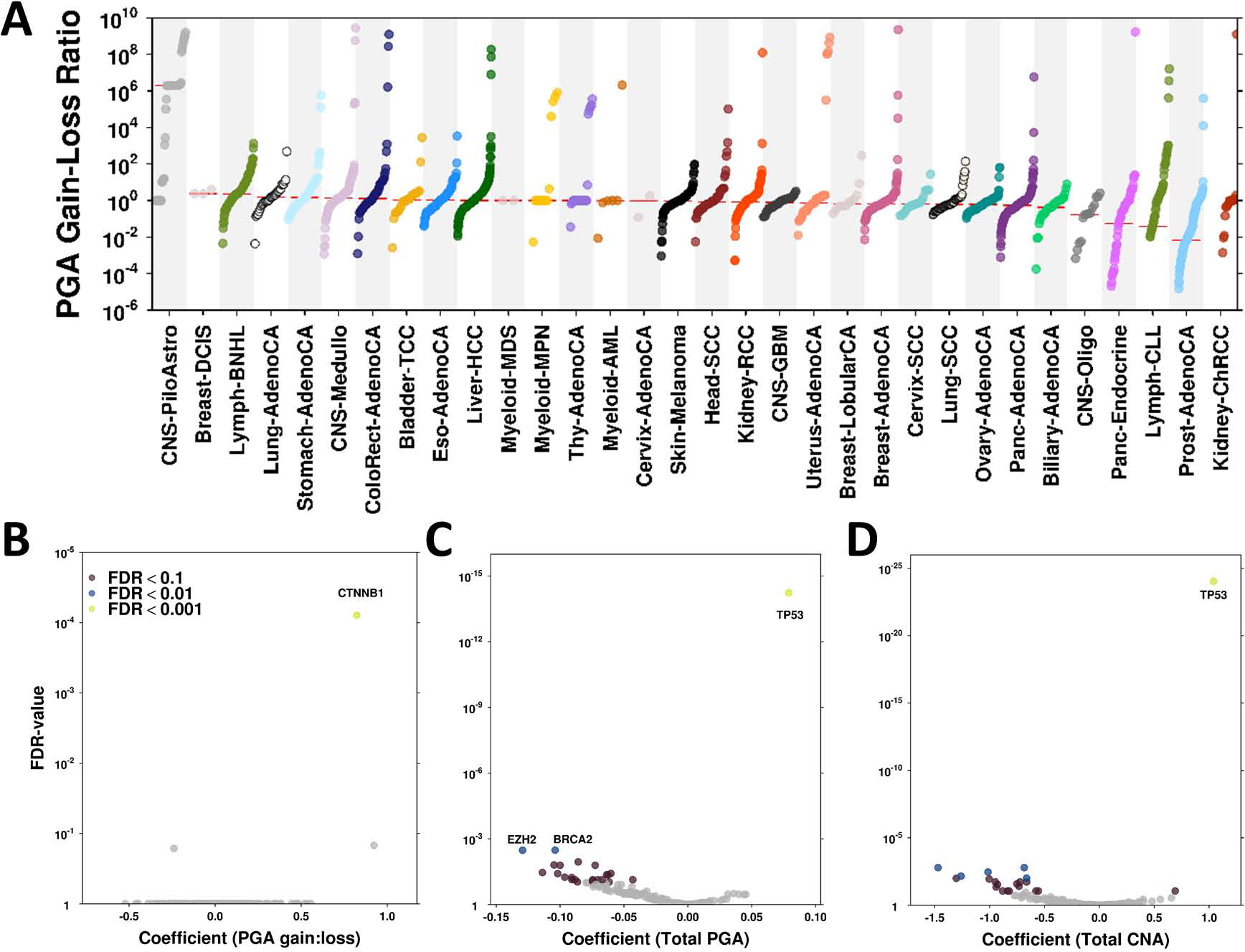
Distinct SNVs are associated with a number of SCNA metrics. A) Ratio of PGA attributed to CN gains vs. losses (autosomes only) differed by several folds across tumour types; tumour types are ordered according to median PGA gain-to-loss ratio; samples within each tumour type are similarly ordered. On the far left, piloastrocytomas harboured few CN deletions, while on the far right chromophobe renal cell carcinoma presented with few CN amplifications (median ratio = 0). Linear mixed-effects regression models were used to find associations between various SCNA metrics and gene-wise SNV status: B) PGA gain-to-loss ratio (autosomes only), C) total SCNA count and D) total PGA, following FDR.

To identify potential drivers of this heterogeneity in SCNAs, we again employed mixed effects modeling. Here, features of SCNA burden were modeled as a function of gene-wise somatic SNV status, considering only those SNVs with predicted functional impact (missense and nonsense mutations single base mutations), and controlling for variables including as patient sex, age and tumour type (**Supplementary Table 3**). Given the relative paucity of recurrent pan-cancer SNVs (<10% of genes contained functional SNVs in ≥1% of tumours), we focused on consensus driver SNVs^28^. We identified a diverse landscape of SNV-SCNA associations across these metrics (**Figure 2B-D, Supplementary Figure 7**). In particular, *TP53* - the most recurrently altered gene (26% of PCAWG tumours harboured at least 1 functional SNV) - was associated with nearly every metric tested, including shorter SCNAs (total or by gains alone), increased number of SCNAs, increased PGA and estimated tumour ploidy. Somatic point mutations in *CTNNB1* was associated with an increased rate of amplifications, relative to deletions (by SCNA count and PGA) as well as with a decrease in overall deletions (both count and PGA) and increased tumour purity. To understand the tumour-type-specificity of these results, we again stratified our analysis and focused on the most well-powered tumour types (100+ tumours) using a similar linear fixed-effects modeling approach (**Supplementary Table 7**). We found minimal recurrence across individual tumour types, with specific genes rarely associated with any one metric in multiple tumour types. The predominant exception to this was *TP53*, associated with metrics of SCNA count (total, gain and loss) and PGA (total and loss) in multiple individual tumour types. Despite the reduced power, these results confirmed the pancancer findings, and highlight a landscape of tumour-type specific effects. These results suggest that in addition to shared biological pathways that increase SCNA mutational density, distinct molecular processes are also at play in different tumour types, and that *TP53* is unique as a strong driver of SCNA burden across many tumour types.

### Drivers of Copy-Neutral Structural Variation

Next, we repeated the above process to assess copy-number neutral somatic structural variants (SVs), in particular translocations and inversions (**Supplementary Table 1**). The mutational density of different SVs were tightly correlated (Spearman’s ρ = 0.81-1.00, p-value < 2.6 × 10^−22^). BRCA-driven tumours (breast, ovarian, and uterine cancers) had high levels of both translocations (TRAs) and inversions (INVs). Alternatively, tumours often thought to be driven by fusion proteins, such as thyroid carcinomas or AML, had fewer total SVs (**Figure 3A, Supplementary Figure 8**).

**Figure 3:**
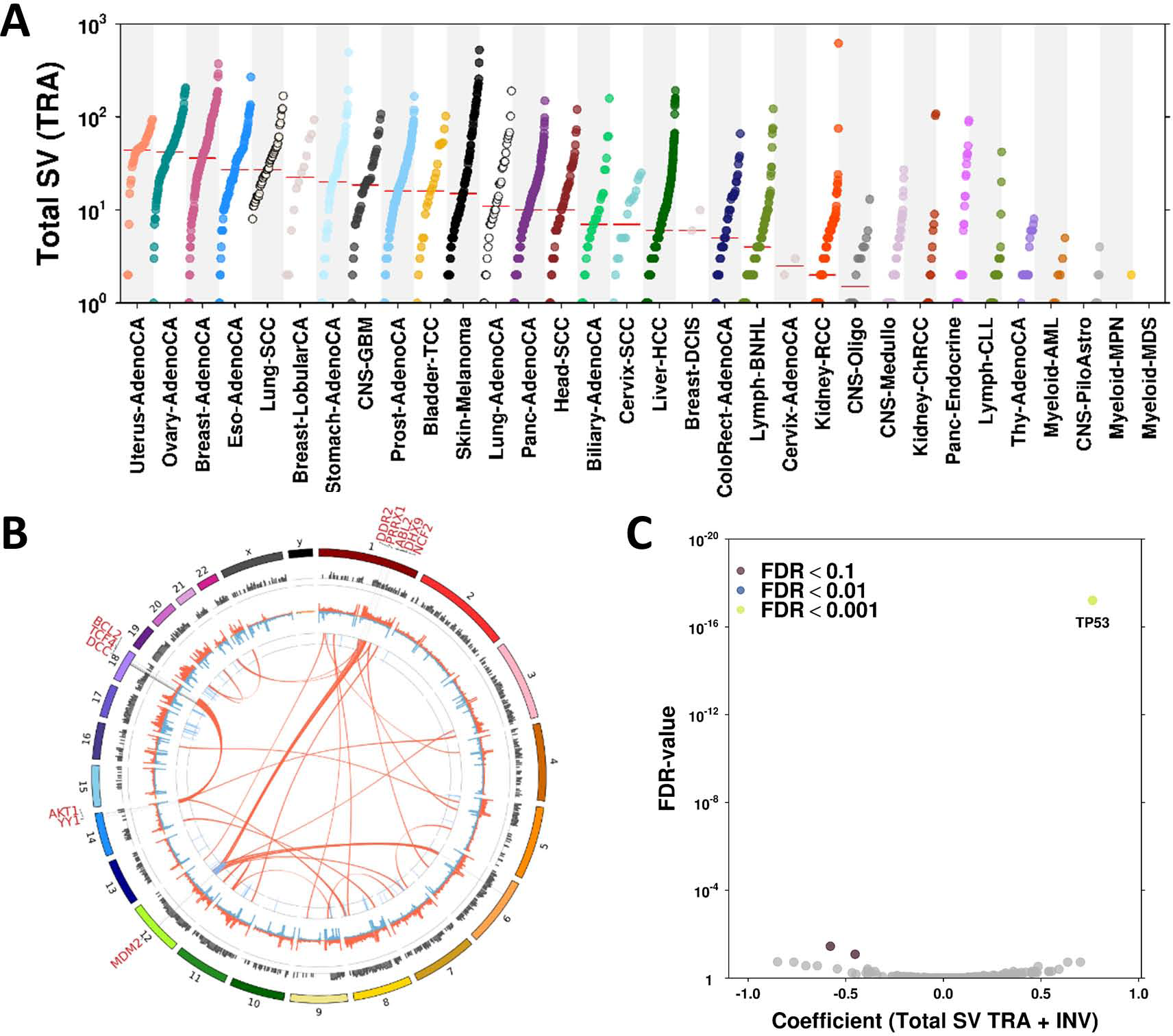
Distinct SNVs are associated with a number of SV metrics. A) The total number of translocations can differ by 2-3 folds across tumour. B) To identify potential patterns in overall SV trend, SVs were first binned and subsequently filtered based on recurrence, this unveiling some interesting patterns in translocations for chromosomes 1, 12, 14 and 18. C) Linear mixed effects modelling identified 9 genes that were statistically significantly associated with translocation count following adjustment for multiple testing.

To understand the spatial structure of these trends, we collapsed SV events into 1 Mbp bins across the genome and visualized those occurring in at least 10 patients (~0.4% recurrence; **Figure 3B**). A total of 149 focal SV hotspots were detected, including two major translocation clusters: one on the q-arm of chromosome 12 (containing *MDM2* and widely distributed across tumour types, however with a predominance in sarcomas) and another between chromosomes 14 and 18 (within regions that include *YY1* on chromosome 14 and *BCL2, SMAD4* and *SMAD7* on chromosome 18) present exclusively in non-Hodgkin’s lymphoma and CLL.

Again, linear mixed-effects modeling was used to identify associations between driver genes containing somatic SNVs and metrics of SV mutational density (**Figure 3C, Supplementary Figure 9**). As expected, results were highly correlated across these density metrics, with somatic point mutations in *TP53* significantly associated with an increase in all SV types (FDR < 0.01). Additionally, point mutations in *CTNNB1, PIK3CA* and *EZH2* were associated with moderately reduced SV density (attributed primarily to inversions; FDR < 0.1). In individual tumour types, *CTNNB1* and *PIK3CA* demonstrated moderate associations with reduced SV count in hepatocellular carcinoma (HCC) and breast adenocarcinoma respectively, while *TP53* showed significant positive associations with total SV count in both tumour types (**Supplementary Table 7, Supplementary Figure 10**). In HCC, point mutations in *TP53* were associated with an increase in inversions (FDR < 0.01); HCC patients with a functional SNV in *TP53* had an average of 1.75x more inversions than those without. Thus point-mutations in multiple genes are associated with an increase in the rate of **somatic SVs.**

### Kataegis

To determine if these broad trends in global mutation burden are mirrored at the level of localized mutational hotspots, we focused on kataegis: localized hypermutation of somatic SNVs^29^. Kataegis can be caused by AID or APOBEC-mediated events^30^, although may also refer to hypermutation due to environmental factors such as UV^31^, and has been described in a number of tumour types^12,29,31-34^. Kataegis is an important signature of genomic instability, independent of other somatic variants^35,36^. To quantify kataegis, we created SeqKat: an open-source tool to predict kataegis from paired tumour and normal WGS *via* a sliding window approach. SeqKat tests deviation of observed somatic SNV trinucleotide content and inter-mutational distance from that expected by chance, after adjusting for the effects of trinucleotide signature and mutation rate (**Supplementary Figure. 11-12**). The resulting kataegic score estimates the magnitude of the event, and accounts for features such as deviation from expected C/T base change frequency and expected inter-SNV distance within each window.

We applied SeqKat to all 2,459 PCAWG tumours with consensus SNV calls and complete clinical annotation and detected 84,218 hypermutation events, of which 97.3% were classified as kataegis (enriched for C>T or C>G mutations in TpCpN trinucleotides). Overall 54% of patients had at least one kataegic event (1,327/2,459 patients; **Supplementary Table 1**), but different tumour types varied dramatically in their number and magnitude. For example, bladder carcinomas had the highest median rate of kataegis, with a median 22 events per tumour, while tumour types had a median of zero kataegic events per tumour (**Figure 4A**). These events also differed dramatically in their scores (length and enrichment for trinucleotide features), with non-Hodgkin’s lymphomas and squamous carcinomas of the lung showing the strongest events (**Figure 4B**). Again, these differences were dramatic, with median event size and score differing by orders of magnitude across tumour types. This heterogeneity among tumour types was similarly reflected within tumour types, with colorectal adenocarcinomas showing large differences in frequency of events (median = 1, SD = 2189) and non-Hodgkin’s B-Cell lymphomas showing the highest variance among in kataegis score (median = 3.2 × 10^5^, SD = 1.15 × 10^6^; **Supplementary Table 2**). Individual events could be remarkably strong, with massive enrichment of the classic APOBEC-associated TCX mutations in some breast tumours (**Figure 4C, Supplementary Figure 13A**) or other, non-APOBEC mediated TCX mutations (**Supplementary Figure 13B-D**). This heterogeneity confirms previous trends in kataegic incidence^11,12,29,32,35,37,38^ and expands them to dozens of additional tumour types, while describing surprising differences in the strength and extent of individual events.

**Figure 4:**
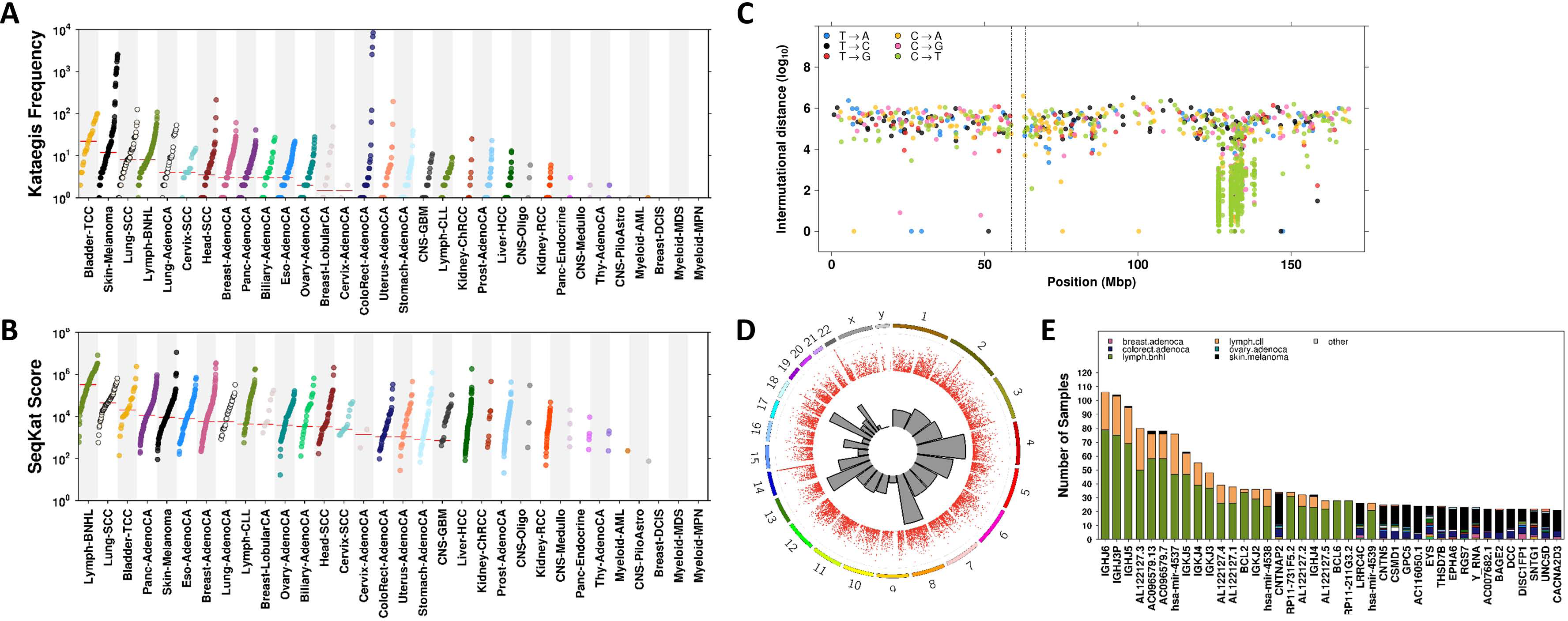
Summary of kataegic events. A) Frequency of kataegis events or B) maximum SeqKat score per patient, stratified by tumour type. C) Rainfall plot depicting kataegis event on chromosome 6 for a single patient with adenocarcinoma of the breast. D) Circos plot displaying the chromosomal distribution of kataegic events in the pan-cancer cohort. E) Top frequent kataegic genes.

To better understand why kataegic events afflict some tumours and regions of the genome more than others, we integrated the presence and location of kataegic events with translocations and RNA-seq data. Chromosomes 4, 5, 8, 18 were enriched for kataegic events relative to chance expectations (q < 0.01; **Figure 4D**). There were 3,479 genes affected by kataegic events in at least 2 patients, ranging from a single tumour type to 11 separate tumour types (**Figure 4E**). *CNTNAP2* harbored the highest number of total kataegic events (n = 304) across 34 patients in different tumour types. This gene, along with 3 other kataegis-enriched genes (*LRRC4C, CNTN5*, and *CSMD1*), showed significant associations with mutational signatures 7a-d (FDR < 0.01; **Supplementary Table 8**), signatures with the characteristic C>T mutations in a TCX context^39^. We observed 7 kataegic hotspots within the genes *IGH, IGK, BCL2, BCL6* and *MYC*, exclusively within lymphomas, consistent with previous studies^40,41^. These genes are either targets for somatic hypermutation (SMG) or aberrant somatic hypermutation (aSHM) in B-Cells^40^. AID and APOBEC editing deaminases play an important role in the initiation of hypermutation and recombination of immunoglobulin genes in B-Cells, which are essential processes for the recognition and disposal of pathogens^42^, explaining the high kataegic rate of *IGK* and *IGH* genes. During that process however, AID has been shown to aberrantly target oncogenes and tumour suppressors such as *BCL6* and *MYC*^41^. Many of these genes are also in close proximity to common fragile sites; while these are typically associated with DSBs and structural variants, fragility may also play a role in hypermutation^43,44^. We assessed the kataegic rate for 56,827 genes, normalizing for gene length, and identified 451 kataegic enriched genes that are potential targets of aSHM in lymphoma (**Supplementary Table 9**).

Aberrant processing by AID also leads to the introduction of translocations *via* induction of double-stranded breaks required by the repair process^45^. We examined the translocation breakpoints in lymphoma patients and found that many of these breakpoints overlapped with kataegic hotspots (**Figure 5A**). One example is the *MYC-IGH* (chr8-chr14) translocation that classically identifies Burkitt’s lymphoma^46^: patients with this event had an enrichment of kataegis events around the translocation breakpoints (10/14 patients). Translocated *MYC* has a consistently lower kataegis score and mutation frequency compared to translocated *IGH*, suggesting that *MYC* kataegic events occurred after the translocation, while under regulation by Ig regulatory elements, as suggested previously^47^. We also assessed the effect of kataegis on *MYC* transcription; in B-cell non-Hodgkin’s lymphomas, kataegic *MYC* had significantly higher mRNA abundance compared to wild-type *MYC* (**Figure 5B**). This form of MYC deregulation has been described previously in B-cell lymphoma cell-lines and can be caused by complex insertional rearrangements, three way recombinations of *MYC-IGH-BCL2* and *IGH-MYC* fusions^48,49^. The presence of Ig transcriptional enhancers in such translocations leads to deregulation of MYC^50^. Similarly, mRNA abundance of most AID/APOBEC family emembers was significantly higher in kataegic samples than non-kataegic ones (**Supplementary Figure 14**), including the transcription factor *PAX5*, which is involved in upregulation of AID^51^.

**Figure 5:**
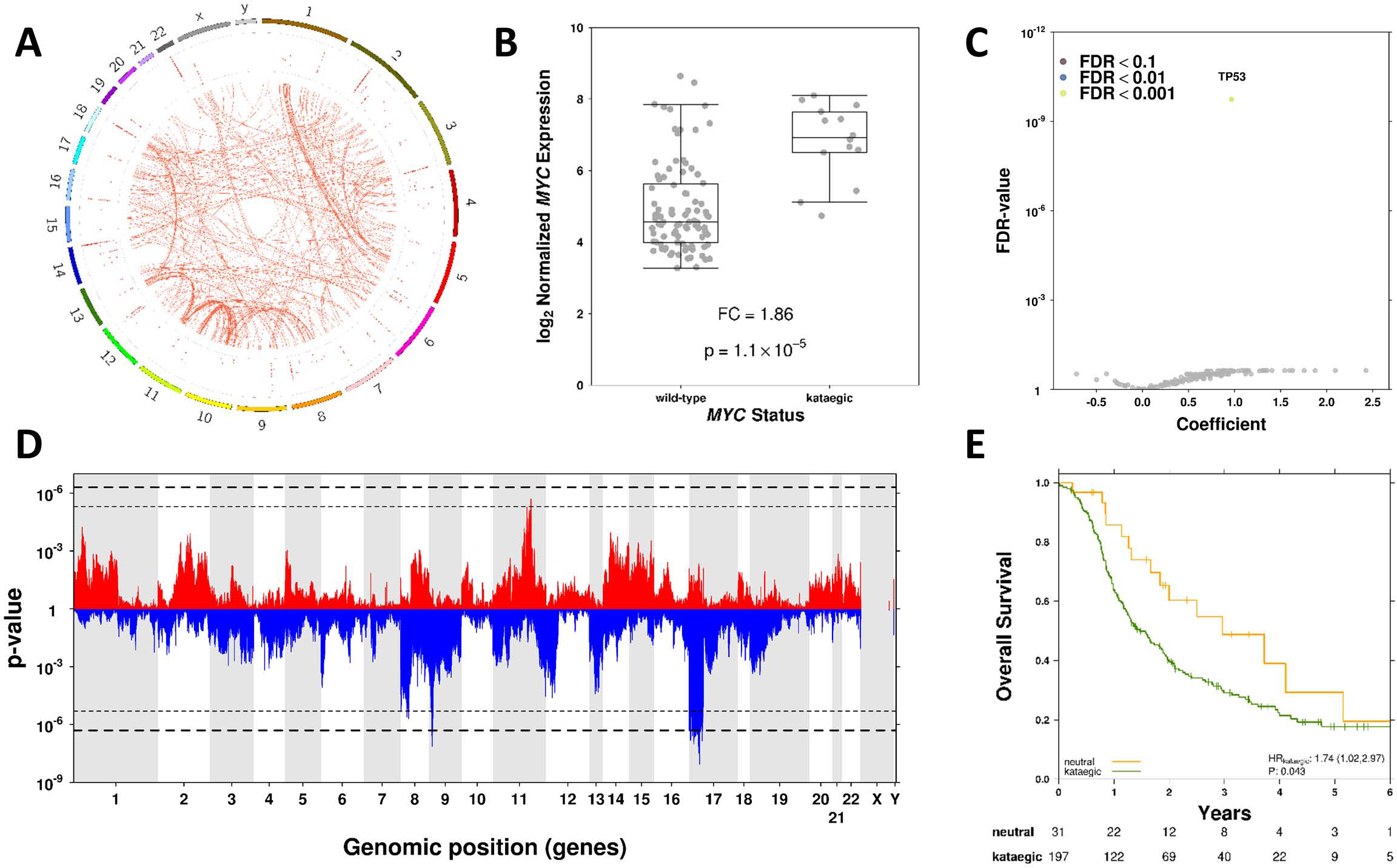
Specific SNVs and SCNAs are associated with an increased in kataegic rate. A) Chromosomal distribution of translocations (centre) in the lymphoma cohort, aligned to recurrent kataegis events (middle layer). B) RNA abundance of *MYC* was significantly higher in non-Hodgkin’s lymphomas patients demonstrating kataegis within this gene than those without. C) Volcano plot demonstrating the results of the mixed effects models using gene-wise SNVs in known driver genes to predict the presence of any kataegis event; a single gene (*TP53*) was significantly associated (FDR < 0.01) with the presence of at least one kataegic event. D) Similarly, p-values for each gene-wise SCNA to predict binary kataegis status were plotted to identify larger, multi-gene events; top (red) bars indicate p-values for gene-wise amplifications while bottom (blue) indicates deletions; dashed black lines indicate Bonferroni-adjusted thresholds (q = 0.1 or 0.01). E) Kaplan-Meier plot demonstrating the overall survival differences between kataegic and kataegis-free patients in pancreatic adenocarcinoma, adjusted for patient sex, age and IGHV status.

We then assessed the relationship between individual gene-specific mutations and different measures of kataegis in each tumour to evaluate its causes and consequences. We again used linear mixed effects modelling to associate the mutation status of each gene with its effect on kataegis, with adjustments for patient sex, patient age, average tumour ploidy (when evaluating associations with SCNA status) and fitting tumour type as a random effect with variable intercepts; both ternary SCNA and somatic SNV status were assessed for each gene (**Supplementary Table 3, Supplementary Figure 15**). Somatic point mutations in *TP53* were remarkably predictive of kataegis (FDR = 1.81 × 10^−10^, effect size = 0.97; **Figure 5C**): tumours harbouring a *TP53* mutation are 2.6x more likely to have at least one kataegic event then those without (**Supplementary Figure 16**). Alternatively, SNVs in numerous genes, including *ERCC6*, were associated with an increase in number of kataegic events (**Supplementary Figure 15A**). When evaluating the impact of gene-wise SCNAs on kataegis, 151 genes contained CN deletions that were associated with both existence and strength of a kataegic event, while 44 genes with CN amplifications were associated only with strength of kataegis event; no genes were associated with the frequency of kataegis events (q < 0.01; **Figure 5D, Supplementary Figure 15C-D**). For example, 19 genes on the p-arm of chromosome 9, including numerous *IFN* genes, demonstrated CN deletion events that were associated with an increased risk of having at least 1 kataegic event. For example, patients with a deletion in *IFNA5* (the most statistically significant event in the locus) were 3.36x more likely to have at least one kataegic event, with a stronger than average signal. A large number of gene-wise CN events were found to be associated with total number of kataegic events in a tumour-type specific manner (q < 0.01 pancancer, q < 0.1 in individual tumour subsets; **Supplementary Table 7**).

Kataegis status, identified through an expression signature, has recently been shown to be associated with late onset, better prognosis and higher HER2 levels in breast cancer^52^. To further investigate the prognostic role of kataegis, a Cox regression was fit for overall survival between kataegic and kataegis-free patients, adjusting for age and sex for each cancer type. As IGHV status is a strong prognostic factor in CLL^53-55^, this was also included as a covariate (**Supplementary Table 10**). Prior to adjustment, kataegis was prognostic in many tumour types, but afterwards only in pancreatic adenocarcinoma, for which the prognostic value of a kataegis event increased slightly on inclusion of IGHV status (p = 0.043, **Figure 5E**); this association was independent of the number of events, only the presence of one or more kataegic events (**Supplementary Figure 17**). Thus kataegis not only appears to be associated with specific driver architectures, but these manifest in a diverse and complex tumour-specific effect on the clinical landscape of outcome and treatment response.

### Shared & Divergent Drivers of Mutational Processes

Throughout our analyses of different types of mutations, *TP53* was recurrently associated with elevated density of almost all mutational features considered. This is consistent with its central role in cancer biology, and a long associating it with DNA damage. These results led us to consider whether there were molecular determinants or correlates of other mutational processes. For each variant type, associations with mutational signatures derived from single base substitutions (SBS) in trinucleotide context, as defined by PCAWG Mutational Signatures Working Group^39^, were evaluated. Of the 65 trinucleotide SBS signatures defined, 23 were present to any degree in at least 1% of the sample population. SNVs, SCNAs and kataegis events were assessed in a gene-wise fashion (n = 1,722, 19,364 and 27 respectively; **Supplementary Table 8**). Interestingly, *TP53* status was rarely associated with these signatures, with point mutations showing a positive association with signature 7c (FDR < 0.01) and CN amplifications positively associated with signature 41 (q < 0.01). In fact, a single gene was rarely associated with more than a single mutation signature, with exceptions including *CNTN5, CNTNAP2* and *LRRC4C*, all of which demonstrate point mutations and kataegis events associated with SBS1 (age) and SBS7a-c (UV-induced pyrimidine dimers). This emphasizes the diverse and complex relationships between characteristics of mutational profiles and individual genes.

Next, we identified common patterns across the overall mutational profile of cancer (**Figure 6A**). Broadly, the mutational density of any given aspect of a tumour was well-correlated with almost every other (**Figure 6B**): tumours that show elevated amounts of one type of mutation tend to show elevated rates of every other type. The one exception was average SCNA length, where tumours with longer SCNAs tend to have fewer of other types of mutations. This is surprising, and may to some extent reflect reduced accuracy of mutation detection in tetraploid tumours, as noted in other studies^26^. To determine whether or not each mutational process was driven by similar mechanisms, we compared the genes significantly associated with each type of mutational density, and found minimal overlap across models for different variant types (SNVs/Mbp, SCNA, SV, and hypermutation/kataegis metrics).

**Figure 6:**
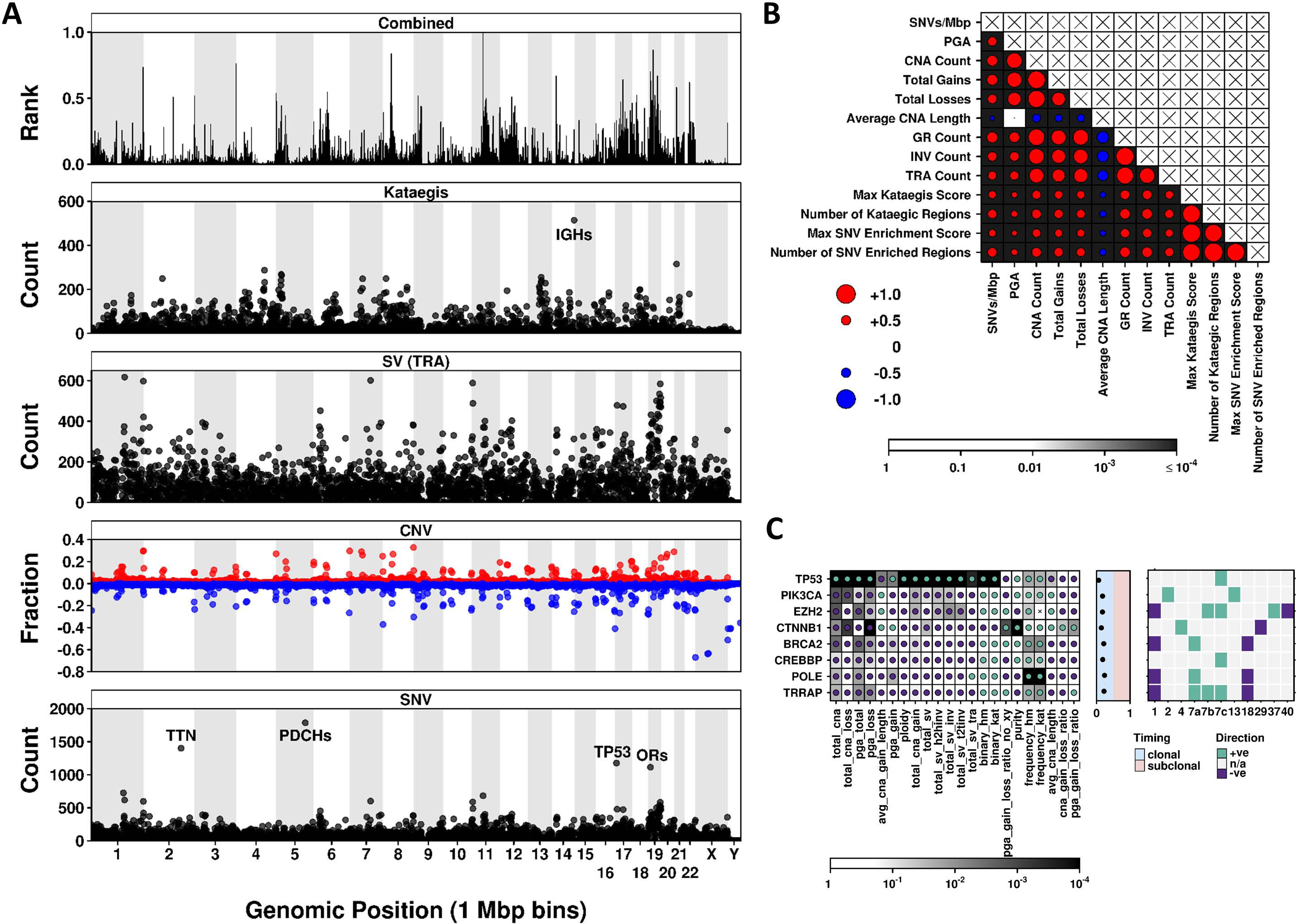
Summary of mutation metrics and key associations. A) Each chromosome was divided into 1 Mbp bins, and the total number of events across patients were calculated for each mutation type, including functional SNV counts, total SCNA counts (divided into gains/losses), total number of transversions, and total number of APOBEC-mediated kataegic events (from bottom to top); counts were then ranked, with the scaled rank product (top) showing mutation enrichment in specific bins. B) Pairwise Spearman’s correlation analyses were performed to assess metric similarity. Dot size indicates the strength of the correlation, with colour indicating direction (red for positive correlation, blue for negative). Background shading indicates the p-value of the correlation. C) Top associated SNVs across the dataset: (left) dot colour represents direction of association (green for positive, purple for negative), while background shading indicates FDR-adjusted p-value; (middle) clonal status of each SNV for the selected genes, dot position indicates the proportion of samples in which this mutation was identified as subclonal; (right) associations with trinucleotide mutation signatures (FDR < 0.01): dot colour indicates direction of associated CN event (red for amplification, blue for deletion) while background shading indicates direction of association (green for positive, purple for negative). For simplicity, all p-values are truncated to 10^−4^.

We then considered the top genes for each mutation density metric (functional SNVs, **Figure 6C**; gene-wise SCNA, **Supplementary Figure 18**). As expected, *TP53* was the most recurrent associated gene, showing a positive association both when mutated by SNVs (associated with many SCNA, SV and kataegis metrics) and when deleted (associated with metrics of kataegis). Numerous genes demonstrated significant associations with metrics of SCNA burden and kataegis, however often with opposite directions. Interestingly, a clear difference between the two mutation types (SCNAs and SNVs) emerged when examining mutational clonality - SCNAs arose predominantly in tumour subclones (with a few exceptions of amplifications arising in the trunk; **Supplementary Figure 18, Supplementary Table 6**), while SNVs were more likely to be classified as clonal (**Figure 6C, Supplementary Table 6**). These results outline both the uniqueness of *TP53* as the only gene associated with almost all types of DNA damage, and the complex, tumour-type specific landscape of mutational drivers revealed by pan-cancer analysis.

## Discussion

One of the most striking results of cancer sequencing studies has been the establishment of dramatic inter-patient variability in the number and nature of somatic SNVs^17^. Some of this variability has been attributed to differences in “mutational signatures”, which reflect the fidelity of a cell’s DNA repair processes and the features of the specific mutagenic insults to which a tumour has been exposed. Here, we focused on identifying the genomic changes associated with these trends in SNV mutation density, and more broadly with multiple classes of mutation density. In particular, we develop new approaches for identifying the associations between mutation-density and candidate driver events, and for detecting kataegic events from whole-genome sequencing data.

These analyses have confirmed a number of previously observed trends, while providing insight into previously unknown oncogenic mechanisms. As expected, tumour types primarily driven by environmental factors, including skin and lung cancers, demonstrated high rates of singlestranded break events while those previously identified as C-class (dominated by copy-number events rather than point mutations)^24^, including ovarian, uterine and breast cancers, demonstrated increased genomic instability, with above average rates of SCNAs and SVs. We quantify how multiple well-characterized driver genes, like *TP53, CTNNB1, PIK3CA* and *MAP3K1* are associated with specific features of the mutational landscape of individual tumours. Further, we confirm the associations of kataegic events with translocations and other complex structural variants.

However these general observations obscure the remarkable divergence of different tumour types. *TP53* is truly an outlier gene, being not only associated with multiple mutational features, but often doing so in individual tumour types. By contrast, many other driver events show Janus-like character. Kataegic events can be associated with either good (*e.g.* GBM) or poor prognostic tumour types (*e.g.* prostate cancer); *DCC* point mutations can be associated with decreased SCNA length (*e.g.* hepatocellular carcinoma) or increased PGA (*e.g.* prostate cancer). Additionally, many of the identified SCNA events identified as associated with increased SNV mutational density and/or kataegis may be confounded by the presence of nearby fragile sites, as these are known to be involved with deletions and other structural events^43,44^, however no comprehensive database yet exists for these data. These divergences highlight the critical importance of appropriate statistical modeling in pan-cancer studies. And, the given the distinctive landscapes and candidate drivers in each tumour-type, these data highlight the ongoing need for large, clinically-homogeneous cohorts with deep WGS to improving our understanding of the mutational hallmarks of individual tumours.

## Methods

### Data assembly and formatting

Clinical annotations were downloaded from the PCAWG data portal on 2016-08 (syn7772065), with survival information obtained from the ICGC data portal on 2017-06-14 (release 25). Samples flagged for removal were excluded and a single sample from each multi-sample donor was selected according to PCAWG recommendations, resulting in 2,583 specimens carried forward for downstream analyses. Consensus SNV calls were obtained on 2016-10-12 (syn7357330) while consensus CNA calls and information on sample purity and ploidy were obtained on 2017-01-25, following the latest PCAWG data release on 2017-01-19 (syn8042905). SV calls were obtained from the 2016-11 release (syn7596712). Callable base files were downloaded on 2017-03-20 (syn8492850). Consensus clonality estimates for SNVs were provided in the 2017-03-25 release (syn8532425) while the latest PCAWG ABSOLUTE calls (2016-11-01) were used to annotate clonal and subclonal CNAs (downloaded from Jamboree on 2017-03-29: /pancan/pcawg11/subclonal_architecture/broad/broad_absolute_on_2660_concensus_bp_11_1 _2016.tar.gz). Candidate driver genes were identified by PCAWG-2,5,9,14 and obtained on 2017-04-22 (syn9757986). Mutation signatures for each patient were downloaded on 2018-05

13(syn11726601)^39^. The RNA-seq expression data was obtained from the 2016-02-12 release (syn5553991). The expression matrix contained upper quantile normalized expression values for 57,821 genes and 2,011 patients.

Consensus SNV calls were filtered such that only functional variants (those predicted to result in either missense or nonsense mutations by Oncotator, as performed by PCAWG-2,5,9,14) were carried forward. Variants were then collapsed to the gene level (n = 18,571 genes), with mutation status further reduced to either present (1) or absent (0) for each patient. These were then filtered to contain only those genes determined to be amongst driver gene candidates (n = 152), identified by PCAWG-2,5,9,14. Similarly, consensus SCNA calls were first filtered based on confidence level (classified by a “star” system, where one star represents poor caller concordance and poor confidence and three stars represent a consensus and high confidence) per patient prior to downstream analysis (more details described here: syn8042880). Only SCNA calls with a star level of two or three were kept. The remaining SCNA calls were adjusted for the estimated ploidy of that sample (based on ploidy data from the latest PCAWG release, obtained on 2017-01-25, data release 2017-01-19, syn8042905), in the event that the sample was predicted to have a whole genome duplication (WGD) with a status of “certain”, and rounded to the nearest whole number. These adjusted calls were referred to as ploidy-adjusted copy number changes and were used in subsequent analyses. SCNAs were annotated to genes using the GENCODE database (v19).

Clonality timing estimates for SCNAs and SNVs were collapsed to form gene by sample matrices, such that for each sample every gene was classified as either clonal, subclonal or not available (indicating either that no variant was present or that timing could not be estimated). A consensus classification was generated across patients (pan-cancer or for the top-powered tumour types) for each gene and event type (CN gain/loss and SNV) using the proportion of patients with a variant that was deemed subclonal in origin (where 0 indicates a clonal event in 100% of patients and 1 a subclonal event; **Supplementary Table 3**).

Sample summary, along with mutation metrics used, are available in **Supplementary Table 1**. All visualizations were made using the BPG package (v5.6.19) for R, with lattice (v0.20-34) and latticeExtra (v0.6-28) packages.

## Data Processing

### Associating overall single-stranded break events with variant status

#### Statistical modeling across the cohort

Uncondensed SNV matrices were first loaded into the R statistical environment (v3.3.1) and overall SNV density was determined for each sample as the total number of single nucleotide variants (SNVs) per callable megabase, ranging from less than 1 to greater than 850 SNVs/Mbp per sample. As this distribution is highly skewed, we defined mutation density for downstream analyses as log10 SNVs/Mbp (determined using the total number of bases with a minimum coverage of 14 or 8 reads in the tumour and normal BAMs respectively; **Supplementary Figure 2A-B**). A total of 2,450 patients had the available callable base data to calculate this metric.

In order to associate specific genomic events with SNVs/Mbp, ternary SCNA status (n = 20,229 genes; n = 1,778/2,450 patients) was used. Genes were first assessed for recurrence to remove those with a SCNA present in less than 1% of the cohort (19,361 genes passed this threshold). For each gene, a mixed effects linear model was applied using the lme4 (v1.1-12) and lmerTest (v2.0-33) packages for R, to explain mutation density across all samples using SCNA status (VS; gain/loss/neutral), sex, age and ploidy with tumour type included as the random effect to set variable intercepts (**Equation 1**). SCNA status, sex and tumour type were treated as factors while age and ploidy were treated as continuous variables. Bonferroni adjustment of the p-values was applied to correct for multiple testing (**Supplementary Table 3**). Tumour purity estimates were not correlated with SNVs/Mbp and were therefore not included as covariates (**Supplementary Figure 2C-F**).

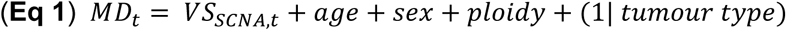

#### Pathway Analysis

For each model term, query lists containing genes significantly associated with SNVs/Mbp (q < 0.01) were generated and separate pathway analyses was performed using gProfileR^56^ (v0.6.1) with default parameters, however with FDR correction for multiple testing and using only the gene ontology (GO) database. Significantly enriched pathways were identified as those with FDR < 0.01 and ordered according to precision (the proportion of term genes present in the query list; **Supplementary Table 4).**

#### Chromosome enrichment

Chromosome enrichment of genes statistically associated with SNVs/Mbp was assessed using genes classified as either protein_coding or processed_transcript in the gencode (v19) database. For each chromosome, a hypergeometric test was used to assess the overlap of significantly associated genes (q < 0.01) and all genes present on that chromosome, from a total pool of all genes (**Supplementary Table 5**).

#### Statistical modeling per tumour type

To increase the power of our analyses, models were run as above on each of the 5 most powered tumour types (those with ≥ 100 samples after filtering). For each tumour type (t), a linear model was applied using variant status (as above), patient age, sex (where applicable) and average tumour ploidy (**Equations 2 and 3**) to identify tumour type specific associations with mutation density. Bonferroni adjustment was applied for each tumour type independently. Analyses were run using ternary SCNA status for collapsed regions (adjacent genes with identical SCNA profiles across each tumour type subset) as the predictor variable (**Supplementary Table 7**).

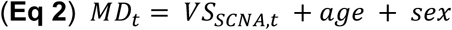

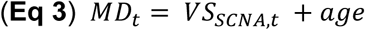

### Associating double-stranded break events with mutation metrics

#### Statistical modeling

Analysis was conducted in the R statistical environment (v3.3.1) for explanation of doublestranded break event metrics (i.e., total SCNA count, PGA, total SV count; full list available in **Supplementary Table 1**). A total of 2,410 samples had the necessary data types for this analysis. Here, driver genes (n = 152) with a functional SNV (described above) in at least 1% of patients were evaluated for associations with these metrics using a similar method to **Eq 1** described above, however without ploidy as a covariate, as all SCNA metrics were previously adjusted for this estimate (to correct for cases of WGD, *etc.).* Here, binary somatic SNV status, considering only functionally impactful variants (missense, nonsense), replaced SCNA status for each gene. Prior to modeling, metrics were transformed as necessary (no transformation for pga_total, purity, cna_gain_loss_ratio, pga_gain_loss_ratio; log10 transformation for total_sv_tra, ploidy; box-cox transform was used for the remainder, using gene-specific lambda values; metrics involving ratios were trimmed to remove extreme values at either end that caused a skew in the distribution, prior to transformation). For response variables showing a non-Gaussian error distribution (ploidy, purity, total_pga and pga_gain), we re-estimated our test statistics using a bootstrap approach (**Eq 4**; tumour types with fewer than 10 samples were combined) and found largely similar results with no optimism bias in the p-values using the linear mixed modeling procedure. For each metric, p-values were adjusted for multiple testing using FDR (rather than Bonferroni, due to the reduced number of tests applied; **Supplementary Table 3**). Models were again repeated for individual tumour types, as described above using a gene-wise approach (modified **Eq 2** and **3** to remove ploidy; **Supplementary Table 7**).

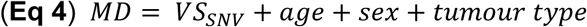

### SeqKat: a tool for assessing hypermutation and kataegis

#### Overview

Genome instability is one of the hallmarks of cancer^57^. A relatively new measure of genome instability is kataegis, which is a pattern of localized substitution hypermutation. Kataegis was first identified in breast cancer^29^ where clusters of C>T and/or C>G mutations were observed in TpCpN trinucleotides on the same strand. Despite the frequent number of studies that have examined kataegis in cancer, its causes and association with other genomic features, there is currently no publicly available bioinformatic tool that can detect and visualize kataegis events per patient using a probabilistic approach. Since kataegis is observed at different rates in different cancer types, a tool that can dynamically optimize kataegis detection per cancer type and assess significance of kataegis events would be of a great use to the scientific community. Here we present **SeqKat**, a novel tool that automates the detection and visualisation of kataegic regions (**Supplementary Figure 11**).

The input to SeqKat is a standard VCF file containing a list of recurrent somatic single nucleotide variants per patient. SeqKat uses a sliding window (of fixed width) approach to test deviation of observed SNV trinucleotide content and inter-mutational distance from expected by chance alone. Additionally, an exact binomial test is performed to test that the proportion of each of the 32 tri-nucleotides within each window is higher than expected. The resulting p-values are then adjusted for multiple hypothesis testing using FDR. Hypermutation and kataegic scores are calculated for each window as follows:

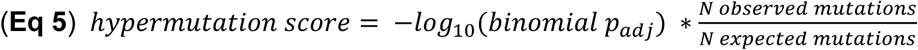

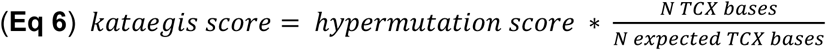

Finally, any statistically significant windows, within an optimized maximum inter-mutation distance, are then combined to obtain regions of hypermutation. Output consists of a text file indicating all potential hypermutated and kataegic regions, their genomic position, and their corresponding hypermutation and kataegic scores.

#### Visualization

The rainfall plot is a comprehensive way of visualizing hypermutation and kataegis events that incorporates both inter-mutational distance and genomic position for each mutation. SeqKat can automatically generate these plots both at the whole genome level and for individual chromosomes (such as only those with a statistically significant event). Hypermutation clusters can be easily recognized using such plots and can be further classified as kataegic events by showing the specific base change composition of the cluster (**Supplementary Figure 13**).

#### Parameter Optimization

Mutation data for 149 samples across 7 different cancer types (ALL, breast cancer, CLL, liver cancer, lung adenocarcinoma, B-cell lymphoma and pancreatic cancer) was downloaded from Alexandrov’s “Signature of mutational processes in human cancer” paper (ftp://ftp.sanger.ac.uk/pub/cancer/AlexandrovEtAl/somatic_mutation_data/)^12^. Data was available as tab delimited files grouped by cancer type. Files were downloaded and converted to BED format per patient. To validate SeqKat, we performed cross validation using these data and tuned the tool’s parameters to maximize prediction performance. Parameters tuned include: 1) Hypermutation score cut-off, used to classify each sliding window as significant 2) Maximum inter-mutation distance cut-off, used to classify significant windows as separate hypermutated events and 3) cut-off for minimum number of SNVs within a single window for it to be classified as hypermutated/kataegic. A 5-fold cross validation was performed and various parameter combinations were run. The combination that maximized the F score across cancer types was selected and used to set defaults (**Supplementary Figure 11D**).

#### Application

SeqKat was applied using the optimized parameters on a cohort of 251 primary whole genome pancreatic cancer samples that are part of the International Cancer Genome Consortium (ICGC). At least one kataegic event was detected in 80% of the cohort. Kataegic samples had an average of four events per sample. Clinical information such as overall survival status (OS), time to OS, grade, age, and sex were obtained for 238 samples. To further investigate the consequence of kataegis on patient overall survival, a Cox regression was fit and overall survival was compared between kataegic and kataegis-free patients adjusting for age and sex. Kataegic patients have significantly poorer prognosis compared to non-kataegis patients (**Supplementary Figure 12**).

#### Download

SeqKat (v0.0.6) is an R package that is currently available in CRAN and can be downloaded from the following link: (https://cran.r-project.org/web/packages/SeqKat/index.html).

#### Assessing Prognostic Role of Kataegis

Clinical information such as overall survival status (OS), time to OS, grade, age, and sex were obtained for 1,704 PCAWG samples. To further investigate the consequence of kataegis on patient overall survival, a Cox regression was fit and overall survival was compared between kataegic and kataegis-free patients adjusting for patient age and sex, as well as IGHV SNV status^53-55^. The analysis was conducted on cancer types that had at least 25 patients with survival and kataegis information available. The test p-values along with the hazard ratios are reported for each cancer type (**Supplementary Table 10**).

### Associating hypermutation and kataegis with mutation metrics

#### Statistical modeling

SeqKat (v0.0.4) was used to identify hypermutation events, classified as either APOBEC-mediated kataegis or not, using the PCAWG consensus SNV calls. SeqKat was run using the default, globally optimized parameters (hypermutation score cut-off = 5, maximum intermutation distance cut-off = 3.2 and minimum SNV count cut-off = 4) and scores were generated as described above. Kataegis events were identified as those hypermutation events with a kataegic score > 0. This identified between 1 and 19,951 kataegic events per sample (mean = median = 1). Where multiple events were called within a single tumour, the event with the highest kataegic score was used to represent the kataegic status of that sample for downstream analyses.

To assess chromosomal enrichment, the genome was split into 1Mbp bins. The expected kataegic rate was calculated by dividing the number of kataegic bins over the total bins in the genome. For each chromosome, the fraction of kataegic bins was calculated and a binomial test was used to test the deviation of observed chromosomal kataegic rate from the expected rate.

For each hypermutation metric (presence of any hypermutation/kataegis event, frequency and score of such events), a mixed effects model was applied as above (**Eq 1**) using either SCNA status (gain/loss/neutral; **Supplementary Table 3**) or gene-wise somatic SNV status of driver genes containing functionally relevant SNVs, again with a 1% recurrence threshold in the dataset (148 genes in 2,563 patients; again, ploidy was not included; **Supplementary Table 3**). Finally, analyses were repeated for each powered tumour type independently (Eq 2 and 3 for SCNAs and modified versions for SNVs, with appropriate distributions, **Supplementary Table 7**).

### Associating specific events with trinucleotide mutation signatures

Analysis was conducted in the R statistical environment (v3.4.0). For each patient, the number of mutations contributing to each trinucleotide profile were obtained and converted to proportions to standardize across patients. Each mutation signature was then modeled as described above using a linear mixed effects model, with SCNA status (gain/loss/neutral, **Eq 1**), presence of SNVs or kataegis events in each gene (modified **Eq 1**, without ploidy), as the independent variable, and controlling for patient sex, age and tumour type. A total of 23 mutation signatures were present to any degree in at least 1% of the cohort. For the independent variables, 1,722, 19,364 and 27 genes had a recurrent event (>1% of samples) related to SNVs, SCNAs or kataegis respectively. Models were run separately for each data type with FDR (SNVs, kataegis) or Bonferroni (SCNAs) adjustment applied to correct for multiple testing (**Supplementary Table 8**).

### Integration across mutation metrics

Mutation enrichment of 1 Mbp bins along the genome was assessed using the rank product of a subset of individual mutation metrics (**Figure 6A**). The large collection of mutation metrics used were compared using pairwise Spearman’s correlations across all available patients (the number of patients differed for each comparison as not all metrics were available for all patients; **Figure 6B**). Furthermore, the top associated SNV or SCNA containing genes from each analysis were compared. Gene-wise functional SNVs were used for associations with DSBs and kataegis metrics, while SCNAs were used to find associations with SSBs and kataegis metrics. For each metric, the top associated genes were selected based on adjusted p-values and magnitude of the coefficient and compared across the different metrics (**Figure 6C, Supplementary Figure 18**).

## Acknowledgment

The authors thank all members of the Boutros lab for technical support and insight commentary. This study was conducted with the support of the Ontario Institute for Cancer Research to PCB through funding provided by the Government of Ontario. Dr. Boutros was supported by a Terry Fox Research Institute New Investigator Award, a CIHR New Investigator Award, by the Canadian Institutes of Health Research grant # SVB-145586, and by Prostate Cancer Canada proudly funded by the Movember Foundation - Grant #RS2014-01.

## Supplementary Figure Legends

**Supplementary Figure 1: Experimental Design.**

**Supplementary Figure 2: Summary of single-stranded break variants**. A) Median somatic SNV frequency across the pan-cancer dataset for each 1 Mbp bin; red points indicate bins encompass known driver genes. B) The distribution of SNVs/Mbp (genome-wide calculation) after log10 transformation. The distribution of SNVs/Mbp according to C) patient sex, D) patient age, E) average tumour ploidy and F) estimated tumour purity coloured to distinguish different tumour types. G) Residuals and predictions were collected from each model; residuals are well distributed.

**Supplementary Figure 3: CN deletions of chromosome 8 associated with SNVs/Mbp in prostate adenocarcinoma**. Model results for individual tumour types. Each panel indicates p-values for each tested gene (with SCNA deletions in at least 1% of the cohort) for a different tumour type. Red line indicates Bonferroni-adjusted threshold for α = 0.1 (gene-wise); blue line indicates same threshold calculated using collapsed CN regions.

**Supplementary Figure 4: CN amplifications of chromosome 1 associated with SNVs/Mbp in medulloblastoma**. Model results for individual tumour types. Each panel indicates p-values for each tested gene (with SCNA amplifications in at least 1% of the cohort) for a different tumour type. Red line indicates Bonferroni-adjusted threshold for α = 0.1 (gene-wise); blue line indicates same threshold calculated using collapsed CN regions.

**Supplementary Figure 5: Several tumours present with notable differences in PGA gain:loss ratio**. The SCNA profiles of five tumour types are shown; CNS-PiloAstro consistently has high PGA gain:loss while Prost-AdenoCA generally has low PGA gain:loss ratio. Within each tumour type, samples are ordered by the PGA gain:loss ratio, calculated using autosomes only, as depicted in top plots. Genomic regions are collapsed by gene and organized along the y-axis by their chromosomal locations.

**Supplementary Figure 6: Summary of DSB mutation metrics (SCNA)**. The distribution of 14 SCNA-derived measures of DSB density across tumours. For each metric, tumour types were ranked according to the median. Dot size indicates rank (where the tumour type with the highest median metric is ranked 1). Background shading indicates variance for each metric in log10 space.

**Supplementary Figure 7: Associations between variant genes and SCNA metrics**. The results of linear mixed effects modelling for assessing gene specific associations with various SCNA-derived metrics of DSB density: Average SCNA length: A) total, B) gains, C) losses or D) gain to loss ratio; total SCNA count divided by E) gains only or F) losses only; PGA calculated for G) gains only or H) losses only; I) Estimated tumour ploidy and J) purity. P-values were adjusted for multiple testing using FDR and statistical significance was defined at FDR < 0.1. To ensure appropriateness of these models, the distribution of the residuals, aggregated across all genes, were evaluated; results were similar for all metrics. For example, K) total SCNA count and L) average SCNA length. Rare exceptions were further contrasted using an alternate method, for example M) distribution of residuals generated using **Eq 1** for PGA by gains and N) correlation of p-values from 2 separate methods; red points indicate results for known driver genes. O) Estimated tumour ploidy demonstrated a non-Gaussian distribution with moderate deviation of p-values between the two methods.

**Supplementary Figure 8: Summary of DSB mutation metrics (SV)**. The distribution of 5 SV-derived measures of DSB density across tumours. For each metric, tumour types were ranked according to the median. Dot size indicates rank (where the tumour type with the highest median metric is ranked 1). Background shading indicates variance for each metric in log10 space.

**Supplementary Figure 9: Associations between variant genes and SV metrics**. Model results for assessing SNV associations with SV-derived metrics of DSB density; volcano plot demonstrating coefficient and FDR-adjusted p-values for known driver genes, and histogram of the residuals, aggregated across all genes, for: A) total SV count, B) total number of inversions and inversions subdivided by C) head to head and D) tail to tail inversions.

**Supplementary Figure 10: Point mutations in *TP53* are associated with SV burden**. Genes found to be associated with metrics of SV burden in a pan-cancer analysis were further assessed in individual tumour types. Four genes were determined to be significantly associated with either A) total SV burden (translocations and inversions) or B) total inversions, at the pancancer level but rarely in individual tumour types. Dot sizes represent the coefficient from the mixed-effects models and background shading denotes the FDR-adjusted p-values. C) Tumours with a functional SNV in *TP53* show elevated SV inversion rates in a pan-cancer multivariate analysis (left boxplot) and in hepatocellular carcinoma (HCC); violin plots stratified by SNV status; coloured violins reflect statistical mixed effects model, FDR < 0.1. The tumour types selected for subgroup analysis are those with the largest sample number (breast adenocarcinoma, pancreatic adenocarcinoma, HCC, melanoma, ovarian and prostate adenocarcinomas).

**Supplementary Figure 11: SeqKat algorithm development**. A) Algorithm workflow highlighting the tuned parameters. B) Overlapping significant windows are stitched together to establish kataegic boundaries. C) Distribution of tumour types used for parameter tuning. D) Performance results following parameter tuning using cross validation. E) Kataegic events can be visualized using rainfall plots; each event has an associated hypermutation and kataegic score.

**Supplementary Figure 12: Application of SeqKat in pancreatic cancer**. Kaplan-Meier plot demonstrating the overall survival differences between kataegic and kataegis-free patients in a pancreatic patient cohort.

**Supplementary Figure 13: Kataegis in numerous tumour types**. Kataegic events were detected in numerous tumour types, including previously described types such as A) breast adenocarcinoma (e.g. chromosome 8 for a single tumour shown), B) B-Cell lymphoma (e.g. chromosome 22 shown) and C) melanoma, where the majority of variants are C>T (e.g. chromosome 17 shown), but also in tumour types previously undescribed as kataegic, for example D) lung squamous cell carcinoma (chromosome 1 shown).

**Supplementary Figure 14: mRNA expression profiles of APOBEC family genes**. Gene expression across the APOBEC family of proteins, stratified by global kataegis status across 912 tumours (across 24 tumour types) with RNA-seq data; groups were compared using a Student’s t-test. A) *AICDA*, B) *APOBEC3B*, C) *APOBEC3D*, D) *APOBEC3C*, E) *APOBEC3G* and F) *PAX5.*

**Supplementary Figure 15: Associations between variant genes and kataegis metrics**. Mixed effects linear modelling was applied to assessing gene associations with kataegis-based metrics. Gene-wise SNV status (driver genes only) was used to predict A) total count of kataegis events or B) maximum kataegis score as a continuous variable, with FDR-adjustment of the p-values. Similarly, gene-wise SCNA status was used to predict C) total count of kataegis events or D) maximum kataegis score, with Bonferroni adjustment of the p-values due to the increased number of tests performed.

**Supplementary Figure 16: *TP53* status is associated with kataegis events**. The proportion of patients with and without kataegis events is significantly different between patients with and without SNVs in *TP53* (Proportion Test P-value = 5.35 × 10^−5^9) in a pan-cancer setting.

**Supplementary Figure 17: Kataegis is associated with overall survival in pancreatic cancer**. Kaplan-Meier plots demonstrating the overall survival differences between kataegic and kataegis-free patients across different tumour types: CLL stratified by A) occurrence or B) frequency of kataegis events; GBM stratified by C) occurrence or D) frequency of kataegis events; E) OV stratified by binary kataegis status; F) pancreatic adenocarcinoma stratified by frequency of kataegis events.

**Supplementary Figure 18: Summary of mutation metrics**. Top associated SCNAs across the dataset; (left) dot colour represents direction of association (green for positive, purple for negative), while background shading indicates Bonferroni-adjusted p-value; (middle) clonal status of each SCNA type (red = amplification or blue = deletion) for the selected genes, dot position indicates the proportion of samples in which this mutation was identified as subclonal; (right) associations with trinucleotide mutation signatures; only Signature 3 was significantly associated (Bonferroni < 0.01) with the selected genes: dot colour indicates direction of associated CN event (red for amplification, blue for deletion) while background shading indicates direction of association (green for positive, purple for negative). For simplicity, p-values are truncated to 10^−4^.

## Supplementary Table Legends

**Supplementary Table 1: Mutation density metrics**. All available mutation metrics for each patient.

**Supplementary Table 2: Summary of mutation density metrics per tumour type**. Median, standard deviation and interquartile range for each metric, stratified by tumour type; values were determined after removing patients with unknown sex/age.

**Supplementary Table 3: Gene-wise model results**. Results of linear mixed effects models for all SSB (using gene-wise ternary SCNAs), DSB (using functional SNVs in driver genes only) and kataegis (ternary SCNAs and functional SNVs in driver genes) metrics in pan-cancer analyses.

**Supplementary Table NN [to delete]: Collapsed, binary SCNA based model results**. Results of linear mixed effects models for binarized SCNA segments predicting SNVs/Mbp.

**Supplementary Table 4: Pathway analysis for genes associated with SNVs/Mbp**. Genes containing statistically significantly associated SCNAs were utilized for pathway analyses. Statistically significantly enriched pathways are shown.

**Supplementary Table 5: Chromosome Enrichment**. Genes that were found to be significantly associated with SNVs/Mbp (when considering CN gains/losses) were assessed for chromosome enrichment.

**Supplementary Table 6: Consensus of clonal/subclonal classification of gene-wise variants**. The proportion of samples demonstrating a subclonal SCNA or SNV was calculated for each gene (proportion ≤ 0.5 = probable clonal variant).

**Supplementary Table 7: Model application to individual tumour types**. Results of linear models for each powered (n ≥ 100 patients) tumour type with the required data for predicting metrics of SSBs (using gene-wise ternary SCNAs), DSBs (using functional SNVs in driver genes only) or kataegis (ternary SCNAs or functional SNVs in driver genes).

**Supplementary Table 8: Mutation signatures are associated with SNVs, SCNAs and kataegis events**. Results of linear mixed effects models for each variant type (genes containing functional SNVs or kataegis events as well as collapsed SCNA segments) modeled independently.

**Supplementary Table 9: Kataegic Enriched Genes**. Summary of genes that are enriched in kataegic events along with the number of samples and tumour types affected.

**Supplementary Table 10: Kataegis association with overall survival**. Summary of Coxph models testing for overall survival differences between kataegic and kataegis-free patients across tumour types with sample size larger than 25 patients.

